# cAMP bursts control T cell directionality by actin cytoskeleton remodeling

**DOI:** 10.1101/2020.07.06.189365

**Authors:** Morgane Simao, Fabienne Régnier, Sarah Taheraly, Achille Fraisse, Rachida Tacine, Marie Fraudeau, Adam Benabid, Vincent Feuillet, Mireille Lambert, Jérôme Delon, Clotilde Randriamampita

**Affiliations:** Université de Paris, Institut Cochin, INSERM, CNRS, F-75014 PARIS, France; Université de Paris, Institut Jacques Monod, CNRS, F-75013 PARIS, France; Master de Biologie, École Normale Supérieure de Lyon, Université Claude Bernard Lyon I, Université de Lyon, F-69342 Lyon Cedex 07, France; Institut de Recherche Servier, F-78290 CROISSY, France; Institut Pasteur, INSERM, F-75015 PARIS, France; Aix Marseille Université, CIML, INSERM, CNRS, F-13009 MARSEILLE, France

## Abstract

T lymphocyte migration is an essential step to mounting an efficient immune response. The rapid and random motility of these cells which favors their sentinel role is conditioned by chemokines as well as by the physical environment. Morphological changes, underlaid by dynamic actin cytoskeleton remodeling, are observed throughout migration but especially when the cell modifies its trajectory. Using dynamic cell imaging, we investigated the signaling pathways involved in T cell directionality control. We monitored cAMP variation concomitantly with actin distribution upon T lymphocyte migration and highlighted the fact that spontaneous bursts in cAMP starting from the leading edge, are sufficient to promote stable actin redistribution triggering trajectory modification.

## Introduction

Fast and random motility of T lymphocytes is a prerequisite to perform efficient immune surveillance, as these cells need to scan the widest possible area in a short time within secondary lymphoid organs (1). This motility is conditioned by the chemical (mainly chemokines) and physical (structural constraints) parameters specifically found in these confined environments. Even in the absence of physical obstacles, random migration is observed (see(2) for instance), suggesting the existence of cell-intrinsic factors regulating the cell directionality.

T cells stimulated by chemokines lose their round shape within a few minutes, to acquire a clear polarized shape with a front, the lamellipodium, and a rear, the uropod. This asymmetry, required for their migration, is achieved by a rapid modification of their cytoskeleton (3). In fact, chemokine stimulation triggers a rapid increase in polymerized actin (4), especially branched actin which accumulates at the cell front giving the lamellipodium some highly dynamic properties adapted to the research strategy of T lymphocytes. Conversely, stable actin allows us to maintain the structural shape of the cell body (5). During T cell migration, continuous remodeling of the cytoskeleton, such as the actomyosin network, has to take place, especially each time cells modify their trajectory. Indeed, in this case, the lamellipodium must first retract, leading to the transient loss of cell asymmetry before being reestablished along another axis. Calcium has been clearly identified as the stop signal leading to lamellipodium retraction and migration inhibition when T cells encounter an antigen-presenting cell (6, 7). However, the signaling pathway involved in shape changes of chemokine-stimulated T cell during trajectory changes, remains unclear.

The role of cAMP upon migration remains confused depending on the cell types or the experimental conditions. In T lymphocytes, a negative effect of cAMP pathway has been known for a long time and is supported by different studies showing that agents inducing large increases in cAMP levels, such as forskolin, inhibitors of phosphodiesterases or prostaglandin E2, promote cell rounding and migration inhibition (8–11). Interestingly, in other cell types, cAMP seems to play a more complex role in cell migration through its compartmentalization. Indeed, in fibroblasts or epithelial cells, an increase of cAMP-activated protein kinase (PKA) at the leading edge has been reported to promote cell migration (12–14). The development of powerful biosensors makes it possible to measure cAMP (15) at the subcellular level even in small cells such as lymphocytes and with a good temporal resolution, and therefore to revisit the role of cAMP in T cell migration.

In this paper, using dynamic cell imaging, we investigate the signaling pathways involved in trajectory control during T cell migration. We demonstrate that transient spontaneous increases in intracellular cAMP are sufficient to drive T cell actin cytoskeleton reorganization, leading to indiscriminate trajectory modifications.

## Results

### Remodeling of actin cytoskeleton in chemokine-stimulated T lymphocytes

Upon chemokine stimulation, T cells lose their symmetrical shape and become polarized. This modification can be visualized by depositing CEM T cells, a lymphoblastic cell line which expresses CXCR4 (the receptor of the CXCL12chemokine), on a glass coverslip coated with the integrin VCAM-1 and CXCL12. In these conditions, cells randomly migrate at a speed of 4.00 ± 0.22 μm/min (n=39 cells) i.e. much faster than highly adherent cells such as NIH-3T3 fibroblasts (16) (illustrated in Supplementary Movie 1). In order to follow in real time and at the subcellular level cytoskeleton reorganization, T cells were transfected with mCherry-tagged LifeAct, a peptide able to bind to F-Actin. As shown in Fig 1a, polymerized actin is observed mainly at the front of the cell which corresponds to newly polymerized actin, as previously described (4). In order to distinguish this pool from stable actin which constitutes the main pool of F-actin in unstimulated cells, SiRActin (17) (a fluorescent cell-permeable F-Actin binding compound) was used. Cells were incubated for 1h with SiRActin and rinsed so that the newly polymerized actin was not labeled. In these conditions, we clearly observed that, contrary to total polymerized actin, the stable actin network was restricted to the back of the cell behind the nucleus (Fig 1a, Supplementary Movie 2). The distribution of the two actin networks was quantified by drawing a scanline along the antero-posterior axis of the cell (Fig 1a) and then by measuring the ratio of intensities between the front and the back (Fig 1b). A ratio superior to 1 indicates an accumulation at the cell front. A statistical difference was measured between the localization of these two actin networks: total polymerized actin accumulates at the cell front while stable actin mainly accumulates at the back. We next wondered whether the polarization of actin networks could be correlated with a mechanical asymmetry during T cell migration. To answer this question, we used the dynamic Traction Force Microscopy technique (18) which allows the forces developed by the cells upon migration on gels to be measured. T cells are fast-moving cells, therefore they are expected to develop low forces on their substrate. For this reason, we used soft gels of about 700 Pa and measured how T cells were able to displace fluorescent beads embedded within the gel while they migrate. As quantified in Fig 1c and illustrated in Supplementary Fig 1a & 1c, T cells clearly imprint centripetal forces with a maximum intensity at the back of the cells and minimal intensity at the front, as previously observed in neutrophils (19). The intensity of the forces we measured was very low, i.e. 1000 times smaller than what was measured in neutrophils on gels with comparable stiffness (19). We can therefore conclude that, during migration, T lymphocytes adhere mainly, but poorly, where stable actin accumulates.

**Figure 1:**
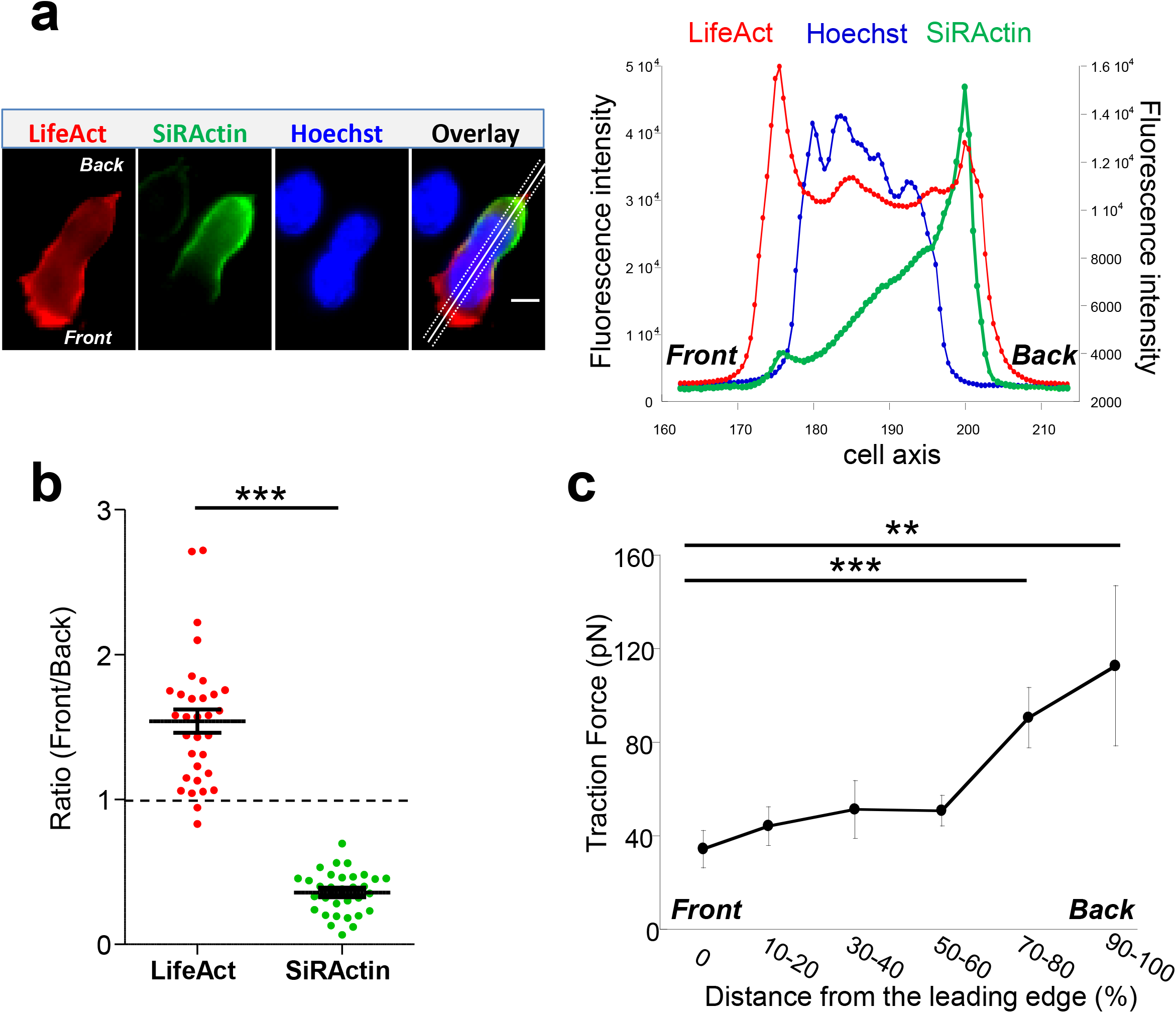
Actin cytoskeleton asymmetry upon chemokine-stimulated T cells. **a**-T cells were transfected with LifeAct-mCherry and labelled with SiRActin and Hoechst. After washing, the cells were deposited on a VCAM-1/CXCL12 coated coverslip. A typical distribution of total polymerized actin (LifeAct), stable actin network (SiRActin) and nucleus (Hoechst) is shown on the left panel and quantified along the scanline displayed on the overlay image (right panel). Images are from Supplementary Movie 2. Scale bar = 10μm. **b**-Using similar scanlines as in **a**, the ratio between front and rear intensities was measured for total polymerized actin and stable actin networks. Mean ± SE (n=31). The values obtained were statistically different (paired t-test, p<0.001, n=32 cells). **c**-The subcellular distribution of forces was measured in CXCL12-stimulated T cells upon migration on approximately 700 Pa gels coated with VCAM-1. The magnitude of these forces was significantly higher at the back of the cells compared to the front. The values of the forces expressed in picoNewton correspond to the mean of forces ± SE measured in 14 different cells. *** p<0.001, **p<0.01 Kruskal-Wallis test.

### Stable actin relocalization upon trajectory modification

The asymmetrical distribution of SiRActin remains stable upon migration. We thus investigated its behavior when cells retract their lamellipodium. This step is specifically required when cells round up and eventually change their direction. The kymograph presented in Fig 2a and the corresponding thumbnails (Supplementary Fig 2a) summarize the different steps: upon migration stable actin remains accumulated at the back of the cell (step 1) & (step 3); the retraction of the lamellipodium is accompanied by the relocalization of stable actin at the front where it rapidly accumulates (step 2) & (step 4). If the cell changes its direction (step 2), the stable actin will migrate entirely to this point which will constitute the new back of the migrating cell, as shown on the kymograph (Fig 2a, red arrow). Conversely, if the cells round up (step 4) (Fig 2a, yellow arrow), the stable actin will progressively redistribute homogeneously all around the cell membrane (step 5). The complete series of images is displayed in Supplementary Movie 3. The distribution of the stable actin during these different steps is quantified in Fig 2b: while, as previously shown in Fig 1b, it is clearly accumulated at the back upon migration (R=0.35 ± 0.01, n=85 cells), a transient accumulation is observed during lamellipodium retraction (R=1.89 ± 0.13, n=53 cells) before it disperses around the membrane (R=0.98 ± 0.04, n=19 cells) when the cell rounds up. A similar relocalization of stable actin was observed when, having developed two lamellipodia, a cell retracts one of them (Supplementary Fig 2b). Lamellipodium retractions were associated with a modification of the cell shape: the cell rounds up before it eventually elongates in another direction. We therefore measured simultaneously over time, the relocalization of SiRActin to the lamellipodium (measure of the ratio front/back along the cell axis) together with the cell roundness as shown in Fig 2c (step (2) of Fig 2a) and measured the delay between the two events (gray arrow). We observed that the accumulation of actin at the front starts at 8.7 ± 4.0 seconds (n=38 cells) before the cells begin to round up, suggesting that the relocalization of the stable actin might drive the retraction of the lamellipodium. Finally, we examined whether the relocalization of stable actin was accompanied by a redistribution of the forces developed by the cells. As shown in Fig 2d and illustrated in Supplementary Fig 1b & 1c, once again the distribution of high intensity forces is similar to that of stable actin: upon retraction, contrary to migratory conditions, centripetal forces at the level of the lamellipodium, reached intensities similar to those observed at the back of the cell. The intensities at the cell front were statistically higher than those observed in migrating cells, while no differences were observed at the back (Fig 2e). Once the cells have rounded, forces can no longer be measured (Supplementary Fig 1d).

**Figure 2:**
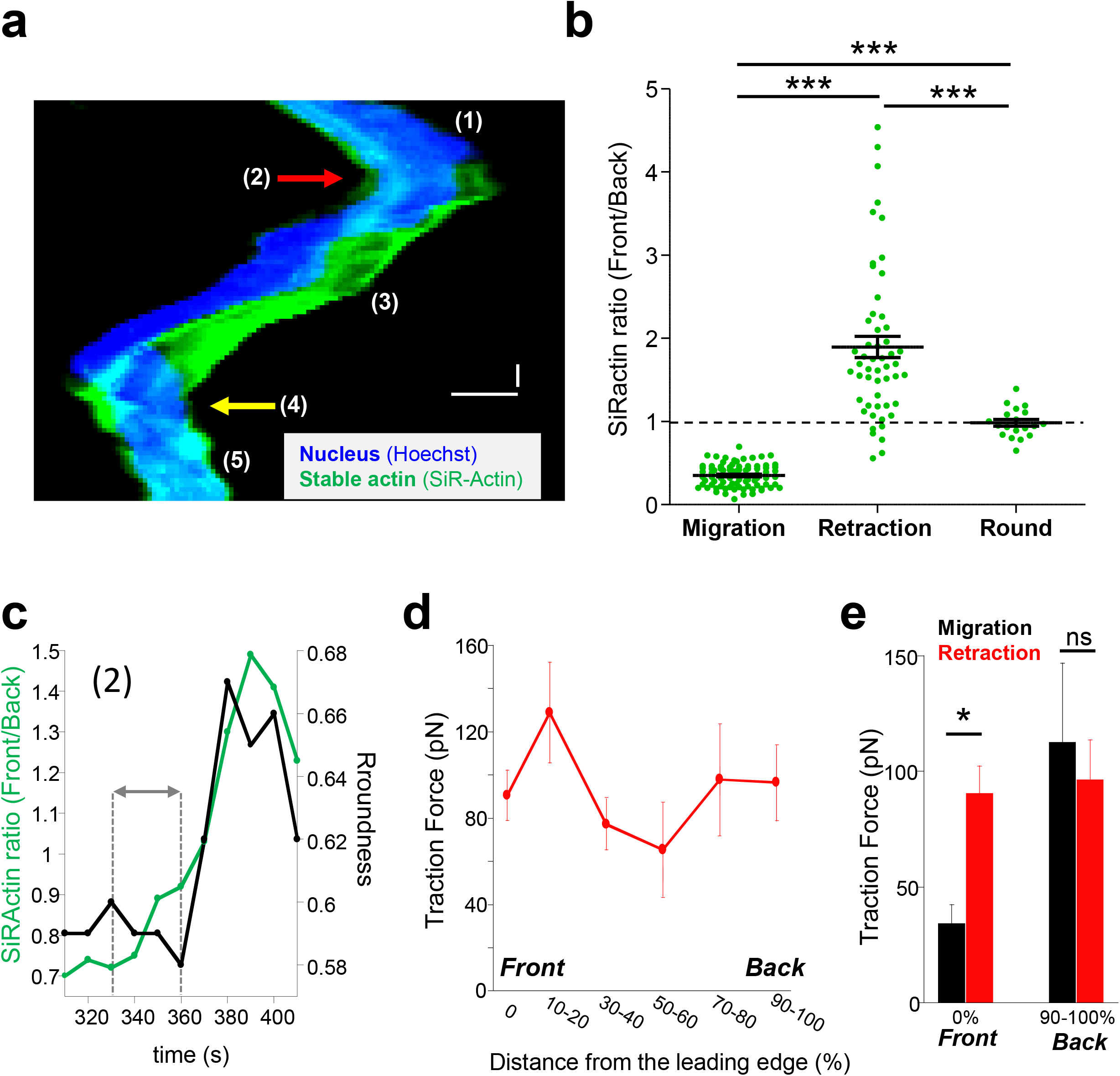
Stable actin relocalization. **a**-After labelling with SiRActin and Hoechst, T cells were deposited on a VCAM-1/CXCL12-coated coverslip and monitored upon migration. A typical example of stable actin distribution and nucleus localization is shown on the kymograph. The x axis corresponds to the average SiRActin intensity along the cell while the y axis corresponds to time. Numbers indicate the different steps: (1) and (3): the cell is migrating. (2), red arrow: it changes its direction. (4), yellow arrow: the cell rounds up. (5): the cell remains round. The associated images are shown in Supplementary Fig 1a and the complete series in Supplementary Movie 3. Horizontal scale bar = 10μm, vertical scale bar = 1min. **b**-The front to back ratio of SiRActin intensities were measured by drawing a scanline along the cell axis of cells upon migration, while the lamellipodium retracted and once the cell had rounded up. Values correspond to the mean ± SE of 95 cells (migration), 53 cells (lamellipodium retraction) and 19 round cells (after retraction). Statistical analysis was performed through a 1way ANOVA test with a Tukey post-test. *** p<0.001. **c**-Cell roundness was measured simultaneously with SiRActin relocalization (ratio Front/Back) during the step (2) (change of direction) for the cell presented in **a**. The delay between the two events was measured (gray arrow). In this example, it corresponds to 30s. **d**-Subcellular distribution of forces was measured in CXCL12-stimulated T lymphocytes upon lamellipodium retraction on approximately 700 Pa gels coated with VCAM-1. The values of the forces expressed in picoNewton correspond to the mean of forces ± SE measured in 8 different cells retracting their lamellipodium. **e**-Comparison of the values of the forces measured in 14 migrating cells (black) or in 8 cells retracting their lamellipodium (red) at their front (0%) or their back (90-100%). * p<0.05 Kruskal-Wallis test.

### cAMP variations upon trajectory modification

We next wondered what the signaling pathway which triggers changes of direction and the simultaneous redistribution of the stable actin might be. Calcium has recently been associated with pausing upon confinement-induced T cell migration (7). Although, in our conditions, calcium transients could sometimes be observed upon migration, they were neither systematic (Supplementary Fig 3a), nor associated with change of direction (Supplementary Fig 3b). We therefore focused on cAMP which has also been described as playing a role during migration(20). We used the very sensitive FRET biosensor, TEpacVV (15, 21) to follow intracellular cAMP levels. As shown on the example presented in Fig 3a and Supplementary Movie 4, cAMP levels remained low upon migration, except at very specific moments when the cell stopped and eventually changed its direction. This can be visualized on the associated kymograph by the red zones corresponding to high cAMP levels. By zooming in on a change of direction (Fig 3a, white dotted rectangle), it appears that the cAMP increase starts at the cell front before invading the whole cell (Fig 3b). By combining cAMP and Ca measurements, we were able to demonstrate that no Ca variations could be detected in cells presenting some cAMP transients upon change of direction (Supplementary Fig 3c). The cellular heterogeneity in cAMP ratio was quantified by drawing scanlines along the antero-posterior axis of migrating cells. The front to back ratios were compared in cells which migrate, retract their lamellipodia or round up. While this ratio is equal to 1.00 ± 0.02 (n=32 cells) upon migration, it increases up to 1.30 ± 0.03 (n=32 cells) when the cells retract their lamellipodia before decreasing back to 1.00 ± 0.04 (n=8 cells) once the cells have rounded up, meaning that lamellipodium retraction is associated with a local increase of cAMP at the cell front (Fig 3c). In our experimental conditions, some cells failed to migrate and went on repetitive elongation/retraction cycles (Fig 3d-f). Interestingly, these cells displayed cAMP oscillations (see Supplementary Fig 4a for two examples) with a very similar period from cell to cell (211.8 ± 11.7 seconds, n=29 cells, Supplementary Fig 4b) and which is very regular for a given cell (Supplementary Fig 4c). These oscillations were associated with morphological changes corresponding to elongation/retraction cycles during which the level of cAMP starts to rise in the lamellipodium before invading the whole cell when it rounds up (Fig 3d & zoom in Fig 3e). The complete series of images is displayed in Supplementary Movie 5. In these cells, the antero-posterior ratio of cAMP was 1.05±0.03 (n=35 cells) upon elongation, increased to 1.38 ± 0.05 (n=35 cells) upon retraction, before decreasing to 0.99 ± 0.03 (n=32 cells) in round cells (Fig 3f,). These values were very similar to those measured in migrating cells (Fig 3c). In order to quantify the coupling between cAMP level and the shape of the cells, the roundness was measured simultaneously with cAMP as in the example presented in Fig 4a (left panel). Clearly, the two parameters oscillate at the same frequency. However, an offset of 50 seconds is necessary in this example to synchronize cAMP levels and roundness (Fig 4a, right panel). A cross-correlation analysis was performed for a series of cells (see methods for details) and reveals that the most significant positive correlation between the two parameters (0.36 ± 0.05, n=17 cells) is obtained with a 40/50s temporal offset (Fig 4b). In other words, this result shows that cells start to round up 40 to 50 sec after cAMP begins to rise.

**Figure 3:**
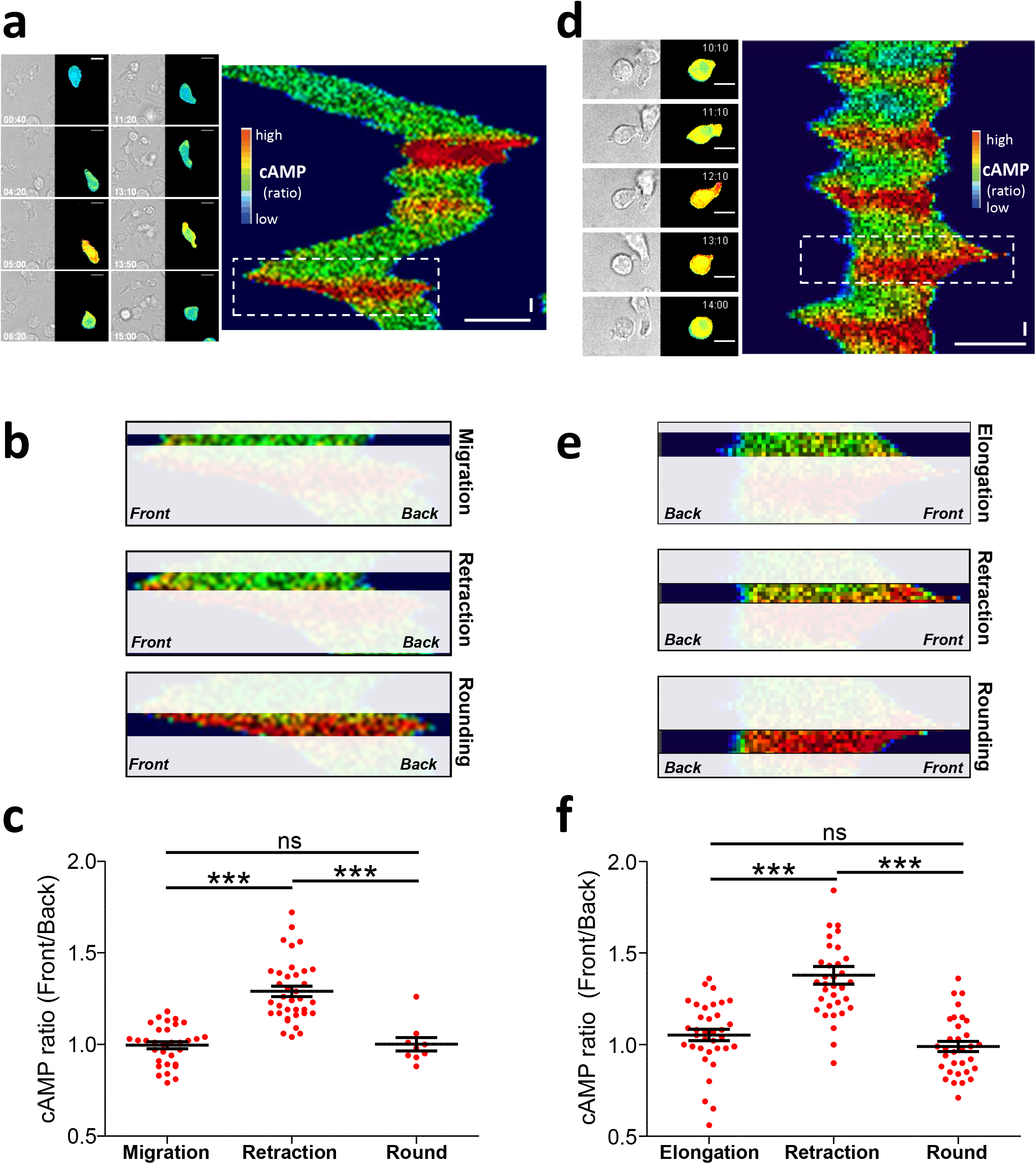
cAMP variations upon migration. **a**-Example of a TEpacVV-transfected T cell migrating on a CXCL12/VCAM-1-coated coverslip. The corresponding kymograph was established along the antero-posterior axis of the cell. The x axis corresponds to the average cAMP level along the cell while the y axis corresponds to time. cAMP levels were coded in false colors. The complete series of images is shown in Supplementary Movie 4. Horizontal scale bar = 10μm, vertical scale bar = 1min. **b**-Zoom of the zone corresponding to the white rectangle in the kymograph presented in (**a**) illustrating that the increase of cAMP starts from the front before invading the whole cell. **c**-Ratios of cAMP level from the front to the back of the cell were measured by drawing scanlines along antero-posterior axis in migrating cells, cells retracting their lamellipodium or after they rounded up. Values correspond to the mean ± SE of 32 events (migration), 36 events (retraction), 8 values (round cells after retraction) from 11 different cells. Statistical analysis was performed through a 1way ANOVA test with a Tukey post-test. *** p<0.0001 **d**-Example of cAMP variations measured in a TEpacVV-transfected T cell displaying elongation/retraction cycles on a CXCL12/VCAM-1-coated coverslip. During recording, the cell presents 5 such cycles as displayed on the kymograph. The complete series of images in Supplementary Movie 5. Horizontal scale bar = 10μm, vertical scale bar = 1min. **e**-Zoom of the zone corresponding to the white rectangle in the kymograph presented in (**d)** illustrating that the increase of cAMP starts at the front before invading the whole cell. **f**-Ratios of front to back cAMP levels measured in cells displaying elongation/retraction cycles. Values correspond to the mean ± SE of 35 values (elongation), 35 values (lamellipodium retraction) and 32 values (round cells after retraction) from 9 different cells. Statistical analysis was performed through a 1way ANOVA test with a Tukey post-test. *** p<0.0001

**Figure 4:**
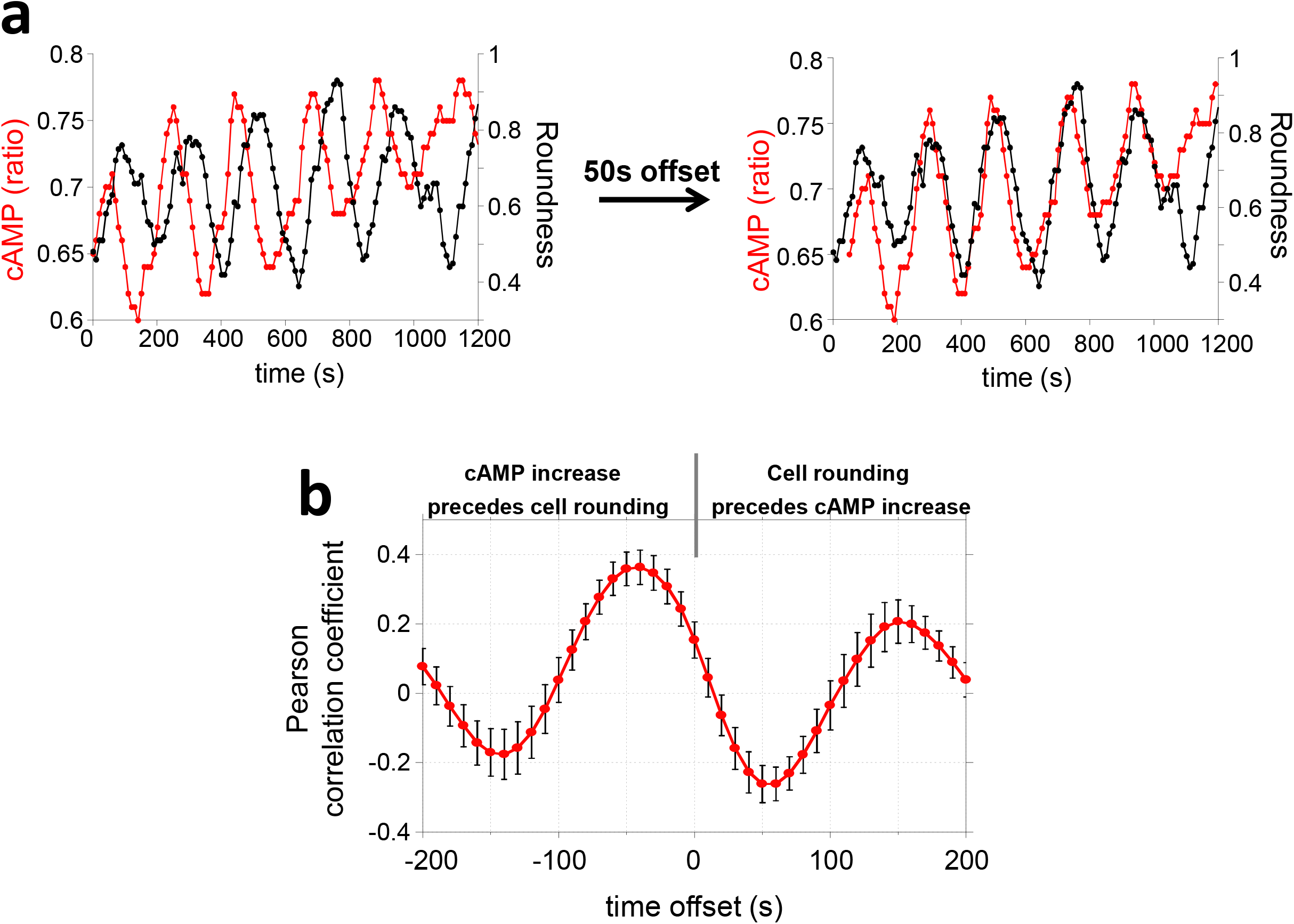
cAMP variations and cell roundness. **a**-Example cAMP variations measured simultaneously with the cell roundness for the cell presented in **Fig 3d**. The shift of cAMP curve by 50 seconds allowed us to synchronize it with the cell roundness curve. **b**-Cross-correlation between cAMP and cell roundness. A negative offset means cAMP increase precedes the cell rounding. Values correspond to the mean ± SE of cross-correlation coefficients measured on 17 cells displaying cAMP oscillations.

### Control of stable actin relocalization by cAMP

To address the direct link between cAMP increase and stable actin recruitment, the two parameters were monitored simultaneously. As shown in the example presented in Fig 5a and in Supplementary Movie 6, a local increase of cAMP can first be observed in the lamellipodium which is followed by a recruitment of stable actin at this position. This observation has been quantified over time, by measuring along a scanline the front/back ratio for cAMP together with the stable actin recruitment (Fig 5a, right panel) and the time lag was measured (gray arrow). The delay between the two events was 40.5 ± 3.6 seconds (n=39 retraction events from 21 different cells, Fig 5b). This result indicates that the local increase in cAMP appears first, followed by the recruitment of stable actin. In order to establish with certainty the causal link between the two events, we artificially generated a local increase in cAMP in the lamellipodium by using a caged form of the nucleotide (DMACM-caged 8-Br-cAMP). The use of this compound allowed us to generate a transient rise in cAMP after illumination at 405nm (22) and to analyze its consequences on the distribution of stable actin, together with cell roundness. Illumination of the leading edge on a 7 μm diameter region induces the recruitment of SiRActin and a lamellipodium retraction (Fig 5c, Supplementary Movie 7). The frequency of retraction upon laser illumination was significantly higher in cells which had been loaded with caged-cAMP compared to control cells (Fig 5d). The retraction events observed in control cells probably correspond to illumination-induced or spontaneous retraction events. After cAMP-induced retraction, cells remain round or form a new lamellipodium in another direction (Supplementary Movie 7). As shown in Fig 5e, the accumulation of stable actin starts 35.2 ± 5.5 seconds (n=23 cells) after cAMP release, while the lamellipodium begins to retract after 48.8 ± 6.5 seconds (n=21 cells). This result demonstrates that a local increase in cAMP is sufficient to induce the recruitment of stable actin and the subsequent retraction of the lamellipodium. Interestingly, by generating an artificial increase in cAMP, the delays measured between the three steps (increase in cAMP / relocalization of stable actin / retraction of the lamellipodium) were very similar to those measured in chemokine-stimulated cells (Fig 5f), suggesting that the signaling cascade is similar in the two configurations.

**Figure 5:**
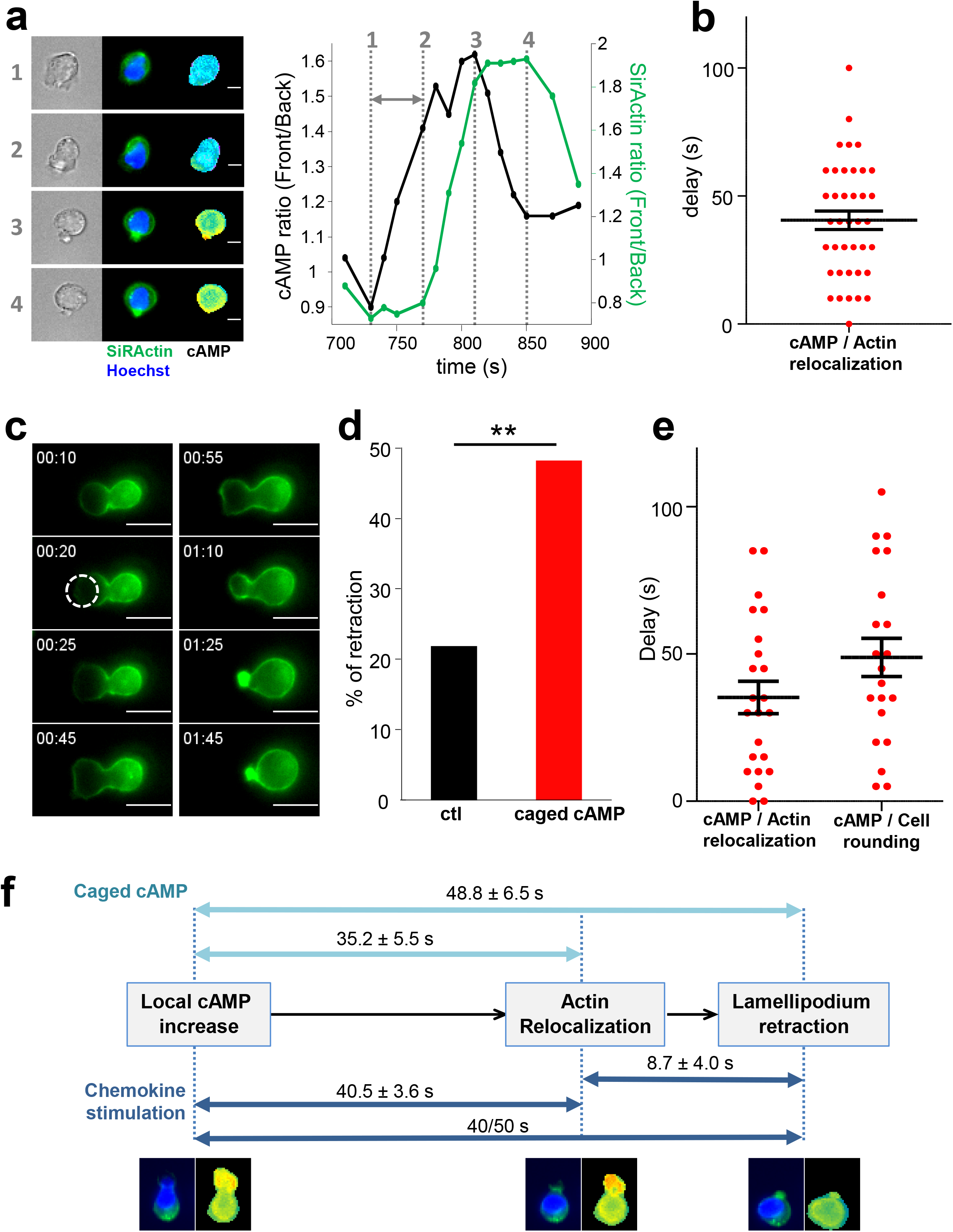
cAMP increase drives stable actin relocalization. **a**-Example of stable actin distribution recorded simultaneously with cAMP variations. The quantification of the two events is shown on the graph and the delay between the two is indicated by the gray arrow. The numbers show the times of the 4 steps. The complete series of images is shown in Supplementary Movie 6. Scale bar = 10μm. **b**-Delays between the beginning of cAMP increase and SiRActin relocalization measured as shown in (**a)**. Values correspond to the mean ± SE of delays corresponding to 39 retraction events from 21 different cells. **c**-Example of rounding up and actin relocalization induced by local release of cAMP after illumination at 405nm. The cell was incubated previously with DMACM-caged 8-Br-cAMP and labeled with SiRActin. The size of the laser spot is indicated by the white circle. The complete series of images is shown in Supplementary Movie 7. Scale bar = 10μm. **d**-The frequency of lamellipodium retraction taking place within 2 minutes after illumination was quantified in cells pre-incubated with DMSO (ctl) or caged cAMP. Analysis of 55 cells for DMSO and 56 for caged-cAMP. ** p<0.01 Chi-2 test. **e**-The delay between cAMP release and stable actin relocalization or increase of the roundness value was measured. For each cell, relocalization of SiRActin was measured over time as well as the roundness. The delay corresponds to the moment at which the values start to increase. Values correspond to the mean ± SE of 23 cells for actin relocalization and 21 for the roundness. **f**-Summary of the three sequential steps leading to cell rounding: local increase in cAMP, recruitment of stable actin at the cell front cell, retraction of the lamellipodium leading to the rounding up of the cells. The delays measured in the different experiments are indicated. Dark blue: observation of migrating cells, light blue: artificial increase of cAMP induced by local photo-release of caged-cAMP.

## Discussion

T cell migration conditions an efficient immune response. The rapid and random displacement of these cells constitutes an important property for an optimization strategy for cognate antigen detection. Although anatomical constraints might impose T cell trajectory, we focus here on the influence of the chemical environment, i.e. chemokine, on T cell migration. We demonstrate that cell intrinsic factors are sufficient to promote random migration upon chemokine stimulation. Indeed, altogether, our results demonstrate that during T cell migration, a pool of actin, visualized with SiRActin, displays an asymmetrical distribution. Surprisingly, this pool is mobile and sets the cell polarity: while it is restricted to the back of the cell upon migration, it is recruited at the lamellipodium upon cell rounding. We have shown that this redistribution is triggered by a rise in cAMP which starts at the cell front before invading the whole cell.

We may wonder, what triggers cAMP bursts observed during T cell migration. The cells displaying repetitive elongation/retraction cycles as observed in some of our experiments might be a good model to address this issue. cAMP oscillations indicate that cells are able to synthetize and degrade cAMP at high frequency (2.5 minutes). Surprisingly, the oscillation period is very similar from cell to cell, which suggests a universal cell-intrinsic cross-talk between adenylate cyclases and phosphodiesterases, the enzymes which respectively synthetize and degrade cAMP. One interesting possibility would be that cell deformation by itself, i.e. membrane stretching, could be the driving force of cAMP bursts. Indeed, the increase in membrane tension generated during migration (23), might drive a cAMP increase, as suggested in other systems (14, 24). In this context, the cAMP-induced recruitment of stable actin would reduce this stretch by retracting the lamellipodium, and therefore inhibit the synthesis of cAMP. In parallel, cAMP increase *via* protein kinase A, one of the main targets of cAMP (25), could activate phosphodiesterases (26) (such as PDE4 highly expressed in T cells (27)) accelerating the cAMP decrease.

The link between cAMP and local recruitment of stable actin is another puzzling observation. As summarized in Fig 5f, a 40-50 second delay is necessary for stable actin to increase at the front after cAMP rise, suggesting that the link between the two events involves a multi-steps signaling cascade. The involvement of PKA is possible, although the nature of its targets remains to be solved. Furthermore, upon lamellipodium retraction, we have observed a restricted zone of stable actin accumulation although the cAMP increase finally invades the whole cell. This suggests that a signal is generated very locally after cAMP increases. An interesting possibility would be the involvement of A Kinase anchoring proteins (AKAP), a family of proteins which would be able to convert the diffusible signal brought by cAMP into a local one like active PKA.

cAMP is generally considered as a messenger which dampens immune response (28). However, this statement must be qualified according to the characteristics of the cAMP increase. Indeed, for T cell activation, although high and sustained cAMP rises have been reported to inhibit TCR signaling such as calcium increase, lck activation or IL2 production (29–33), we have previously shown that T cell adhesion to antigen-presenting cells triggers a transient increase in cAMP which lowers the antigen detection threshold and therefore favors T cell response (22). Concerning migration, similarly, high and sustained cAMP rise triggered by pharmacological drugs, PGE2 or ß-adrenergic receptors stimulation (8–11), are known to inhibit T cell motility. However, our present study highlights that, conversely, transient burst in cAMP, by remodeling the actin cytoskeleton, might favor the exploratory behavior of T cells, a crucial step to mounting an efficient immune response. It might therefore be important to revisit the immunosuppressive effect of cAMP. Spatiotemporal control of cAMP signal is therefore crucial for T cell properties: although sustained rise of cAMP may be inhibitory, transient increase of this messenger may, conversely favor T cell response.

## Acknowledgements

We thank G. Bismuth for helpful discussions and comments on the manuscript, Anna Mularski for helpful advice on Traction Force Microscopy, J.L. Martiel and Q. Tseng for Image J plugins for Traction Force Microscopy and advice, K. Jalink for TEpacVV constructs and the IMAG’IC facility for technical advice on local uncaging. This work was supported by Cochin Institute (PIC Program), Association pour la Recherche contre le Cancer (PJA 20131200379), CNRS, INSERM and Université de Paris. MS was supported by the Ministère de l’Enseignement Supérieur et de la Recherche.

## Author contributions

MS, FR, ST, AF, RT, MF, AB, VF, JD, CR performed experiments and analyzed data. ML helped with the experimental design of Traction Force Microscopy experiments. AF wrote R code for gel rigidity measurements. MS, JD and CR designed experiments and wrote the manuscript.

## Methods

### Cells

CEM T cells were cultured in RPMI 1640, supplemented with 10% FCS, 2mM L-Glutamine, 50 U/ml penicillin and 50 μg/ml streptomycin. When specified, cells were transfected by nucleofection (Amaxa Nucleofactor, Lonza) with 5μg DNA for 5M of cells using the C-016 program. The cells were used the day after nucleofection.

### Reagents

CXCL12 (recombinant human SDF1-α) was purchased from Peprotech (300-28A) and VCAM-1 (CD106 Fc chimera protein) from R&D Systems (862-VC-100). Calcium measurements were performed with Fura-2/AM (Molecular Probes, F1225). DMACM-caged 8-Br-cAMP was purchased from Biolog (D044). Nucleus labeling was performed with Hoechst (Molecular Probes, H1399). F-actin detection was performed by expressing the LifeAct-mCherry construct (gift from Dr A. Benmerah). Stable actin detection was performed with SiRActin (TebuBio, SC001).

### Live imaging acquisition

For migration experiments, glass coverslips were coated with 1μg/ml CXCL12 and 1μg/ml VCAM-1 overnight at 4°C. After rinsing, coverslips were kept in mammalian saline buffer (140 mM NaCl, 5 mM KCl, 1 mM CaCl_2_, 1 mM MgCl_2_, 20 mM HEPES, 11 mM glucose) supplemented with 5% FCS. Cells were deposited on coverslips just before image acquisition started. Live imaging experiments were performed at 37°C with a wide-field Nikon TE2000, equipped with a CMOS camera (ORCA-flash4.0 LT, Hamamatsu). Images were acquired every 10 seconds with Metafluor software.

#### Actin and nucleus

For total polymerized actin detection, cells were transfected with LifeAct-mCherry construct. For stable actin labeling, cells were incubated for 1h with 250nM SiRActin in complete medium at 37°C. After rinsing, cells were deposited on coated coverslips. Distribution of actin was followed by 650nm excitation and 700nm emission (SiRActin) or 560nm excitation and 645nm emission for LifeAct. Nucleus labeling was performed with 4 minutes incubation, using 2μg/ml Hoechst.

#### cAMP measurements

For cAMP measurements, cells were transfected with the most sensitive version of TEpacVV (H187 (21)). TEpacVV was a gift form Dr. K. Jalink (Netherlands Cancer Institute). Experiments were performed 24h after transfection, as previously described(22). Briefly, when cAMP increases, the probe undergoes a conformational change that allows a decrease of energy transfer between a turquoise molecule (excitation 436 nm, emission 470 nm) and two Venus molecules (Exc 500 nm, Em 535 nm) (15); the energy transfer can be measured as a change in FRET (Exc 436 nm, Em 535 nm). Three images were acquired every 10 s: visible, Turquoise channel and FRET channel. The ratio R = Turquoise/FRET, which gives an estimate of cAMP concentration, was calculated with MetaFluor (Roper Scientific) after background subtraction. An increase of this ratio corresponds to an increase in cAMP concentration.

#### Calcium measurements

For calcium experiments, cells were loaded with 500nM Fura-2/AM for 20min at 37°C. Excitation was performed alternatively at 350 and 380 nm and emission recorded at 510 nm. The ratio (Exc 350, Em510 / Exc380, Em510) was calculated with MetaFluor (Roper Scientific) after background subtraction. For combined cAMP and Ca measurements, TEpacVV-transfected cells were loaded only with 200 nM Fura-2/AM in order to minimize the crosstalk between the four fluorescence signals.

### Image analysis

#### Cell roundness

This parameter was quantified with ImageJ software and corresponds to 4xArea/Πx(Major Axis)^2^. It is equal to 1 for a round cell and <1 for an polarized one.

#### Front/Back ratios

With Metamorph software a scanline 6-9 pixels wide along the cell axis was drawn. The front/back ratio was then calculated by dividing the intensity at the front edge to the one measured at the back one.

#### Kymographs

Kymographs have been performed with Metamorph software by drawing a line as wide as the cell along the migration axis.

#### Cross-correlation

Cross-correlation was used to study the correlation between cAMP variations and cell shape (roundness). The Pearson correlation coefficient (ρ) between two time courses was computed as a function of time lag (τ):

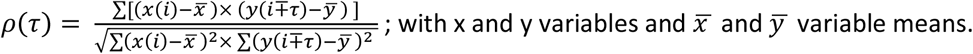

### Local leading edge release of cAMP

CEM were incubated with 20μM DMACM-caged 8-Br-cAMP for 3h at 37°C. SiRActin 250nM was added to the medium during the last hour. Cells were rinsed and deposited on VCAM-1/CXCL12 coated coverslips for 30min at 37°C. The experiment was performed by using an iMIC TILL Photonics microscope equipped with 2 cameras EMCCD (ANDOR Technology) and 60x objective (numerical aperture: 1.49) + 1.5 zoom. Images were acquired every 5s. After 4 image acquisitions, the release of DMACM-caged 8-Br-cAMP was performed with a 405nm laser (Toptica iBAEM 110mV, 1ms illumination, 100% power) by adjusting a 7μm diameter region at the level of the leading edge of a migrating cell. For control experiments, a similar protocol was followed except that the cells were incubated with DMSO for the same duration.

### Traction Force Microscopy

Traction force microscopy experiments were performed with the help of Cell Biomechanics facility of Cochin Institute.

#### Hydrogel preparation

Hydrogels (~700Pa) were prepared with acrylamide (3%, Sigma #A4058), bis-acrylamide (0.3%, Sigma #M1533-25), streptavidin-acrylamide (Invitrogen, S21379) and Flash Red 0.2μm fluorescent beads (Bangs Laboratories, FSFR002). Streptavidin-acrylamide was used at 1/100000 molecular ratio to acrylamide as previously described (34). After activation with TEMED and ammonium persulfate, 11μl of the polymerization mix was added on a non-functionalized 12mm diameter coverslip. A functionalized glass coverslip coated with silane (Sigma, 17-1330-01) was placed on top. Polymerization was performed at room temperature for 30 minutes.

#### Mechanical properties of the polyacrylamide gels

Gels were unmolded by removing the non-functionalized coverslip. We then checked whether the bead distribution on the top surface was suitable for traction forces measurement (~2000 beads per 512×512 pixels field). The Young Modulus was then calculated according to (35). In brief, tungsten carbide spheres with known radius (0.4 and 0.6mm) and density (15630g/l) were deposited on the hydrogel surface. We then measured the gel deformation induced by the bead by acquiring z-images of the fluorescent beads embedded in the gel, focusing on the bottom and the top of the gel with an indentation of 0.2μm. By using ImageJ, we measured the gel height and the collapse distance of the sphere. The Young Modulus was calculated by using a R code based on (35) (available on demand).

#### Functionalization of hydrogel surface

We used the specific biotin-streptavidin binding and anti-Fc/Fc binding to form a sandwich of macromolecules for the functionalization of polyacrylamide gels. All gel surfaces were incubated with 10 μg/ml of a goat anti-human IgG Fc biotinylated antibody (Abcam: ab97223) in PBS-BSA 0.2% overnight at 4°C. Gels were then incubated with 10μg/ml recombinant human VCAM-1/CD106 Fc chimera in PBS-BSA 0.2% for 2 hours at 37°C. We were not able to experimentally assess the VCAM-1 surface density, but theoretically calculated the density of streptavidin molecules on the gel. For this we used the three assumptions enunciated in (34). In brief, 1) the volume of the hydrated gel (with culture medium) that we were able to calculate with the thickness and the coverslip diameter, is approximately 40% bigger than the initial volume of the polymerization mix (36). 2) All the streptavidin-acrylamide molecules within the polymerization mix polymerized within the gel. 3) Biotinylated anti-Fc antibody can access the first 10nm of the gel (10nm is the approximated size of the streptavidin molecule) due to their own size and the size of the pore reported in the literature (37).

We then calculated that the theoretical surface density of streptavidin-acrylamide is 25 molecules / μm^2^.

#### Traction force measurements

Traction force microscopy experiments were performed with 20x objective (numerical aperture 0.75) and 1.5 zoom. CXCL12-stimulated (100ng/ml) cells were deposited on a VCAM-1-coated gel for 30min at 37°C. Transmitted light and corresponding fluorescent images of beads and actin were acquired every 10s using the MetaMorph software.

#### Force image analysis

We first aligned images of the fluorescent beads to correct the drift by using the ImageJ plugin Stack Reg. The forces were calculated by the method described in (38). Basically, the displacement field was calculated by Particule Image Velocimetry (PIV) plugin implemented in ImageJ. The PIV was performed through an iterative process. For each iteration, the displacement was calculated by the normalized correlation coefficient algorithm, so that an individual interrogation window was compared with a larger searching window. Each subsequent iteration took into account the displacement field measured previously. The resulting final grid size for the displacement field was 5.04 x 5.04 μm with more than six beads per interrogation window on average. With the displacement field obtained by PIV analysis, the traction force field was reconstructed by the Fourier transform traction cytometry (FTTC) method(38) with FTTC ImageJ plugin. The regularization parameter was set at 8 x 10^-11^ for all traction force reconstructions.

After this calculation, the forces along the cell body were isolated. The cell length was normalized by establishing that the cell front corresponds to 0% and the back to 100%. When specified (Fig 1d, Fig 2c-d), the forces along the cell axis were pooled.

### Statistics

The statistical tests used for sample comparison are specified in the figure legends. They were performed with GraphPad software or RStudio.

## Supplementary Figure Legends

**Supplementary Figure 1:**
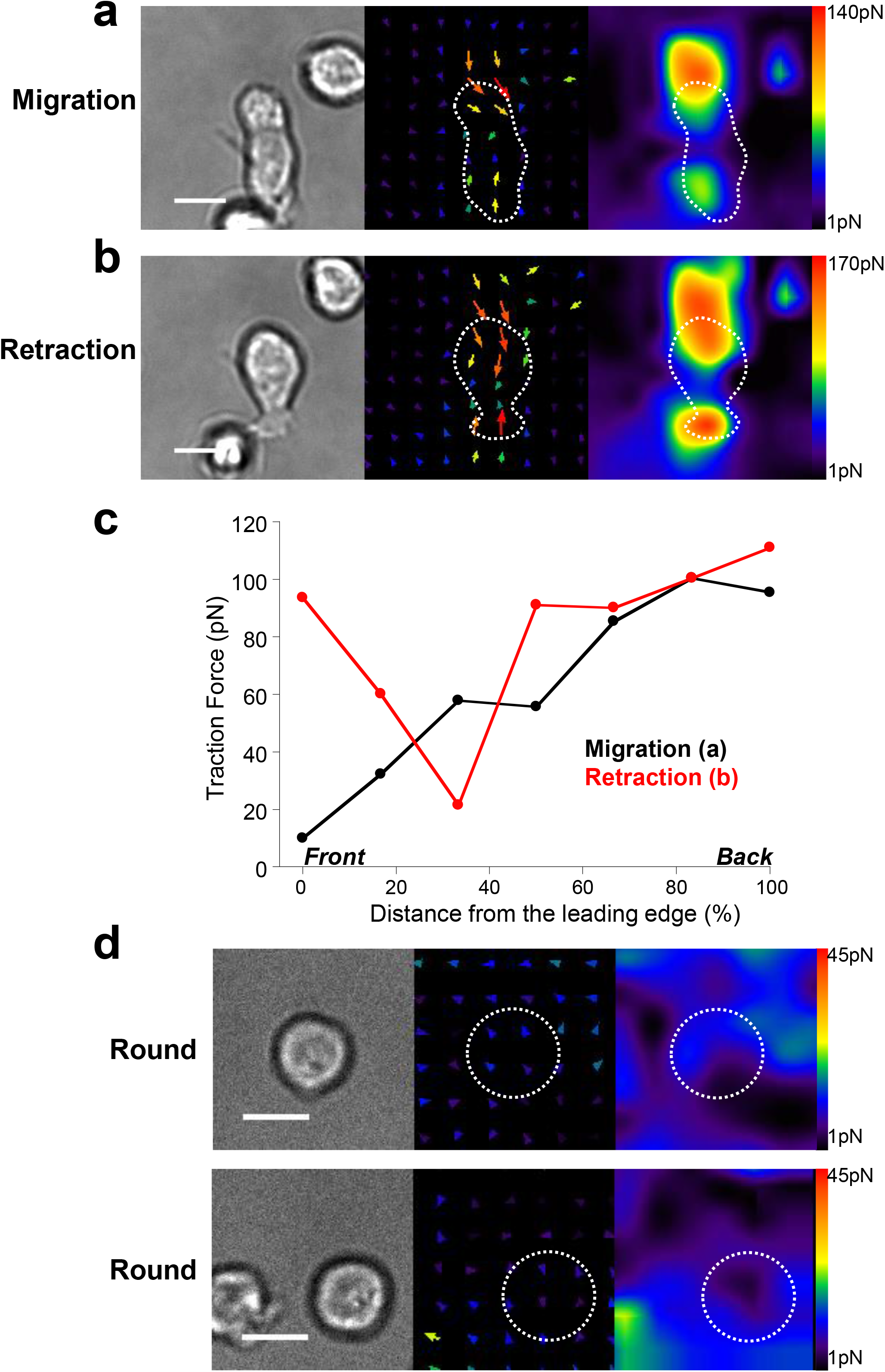
Example of Traction Force Microscopy. **a-** CXCL12-stimulated cell migrating (left panel: transmitted light picture) on VCAM-1 coated gel develops centripetal forces (vectorplot, center panel) whose magnitude is at a maximum at the back (Heatmap, right panel). The cell outline is indicated by a white dotted line. Note that the size of the gel area deformed by the cell is larger than the cell itself. The color code indicates the traction force intensity in picoNewton. **b-** The same cell as in **(a)** is retracting its lamellipodium. Although the forces are still centripetal (vectorplot, center panel), similar intensities can be measured at the back and at the cell front (Heatmap, right panel). As in **(a)** the size of the gel area deformed by the cell is larger than the cell itself. The cell outline is indicated by a white dotted line. **c-** Quantification of the forces along the cell presented in **(a** & **b)** upon migration and retraction of its lamellipodium. d-Two examples of cells which no longer imprint forces on the gel after lamellipodium retraction. Vectorplot in the center and Heatmap on the right. Scale bar = 10μm

**Supplementary Figure 2:**
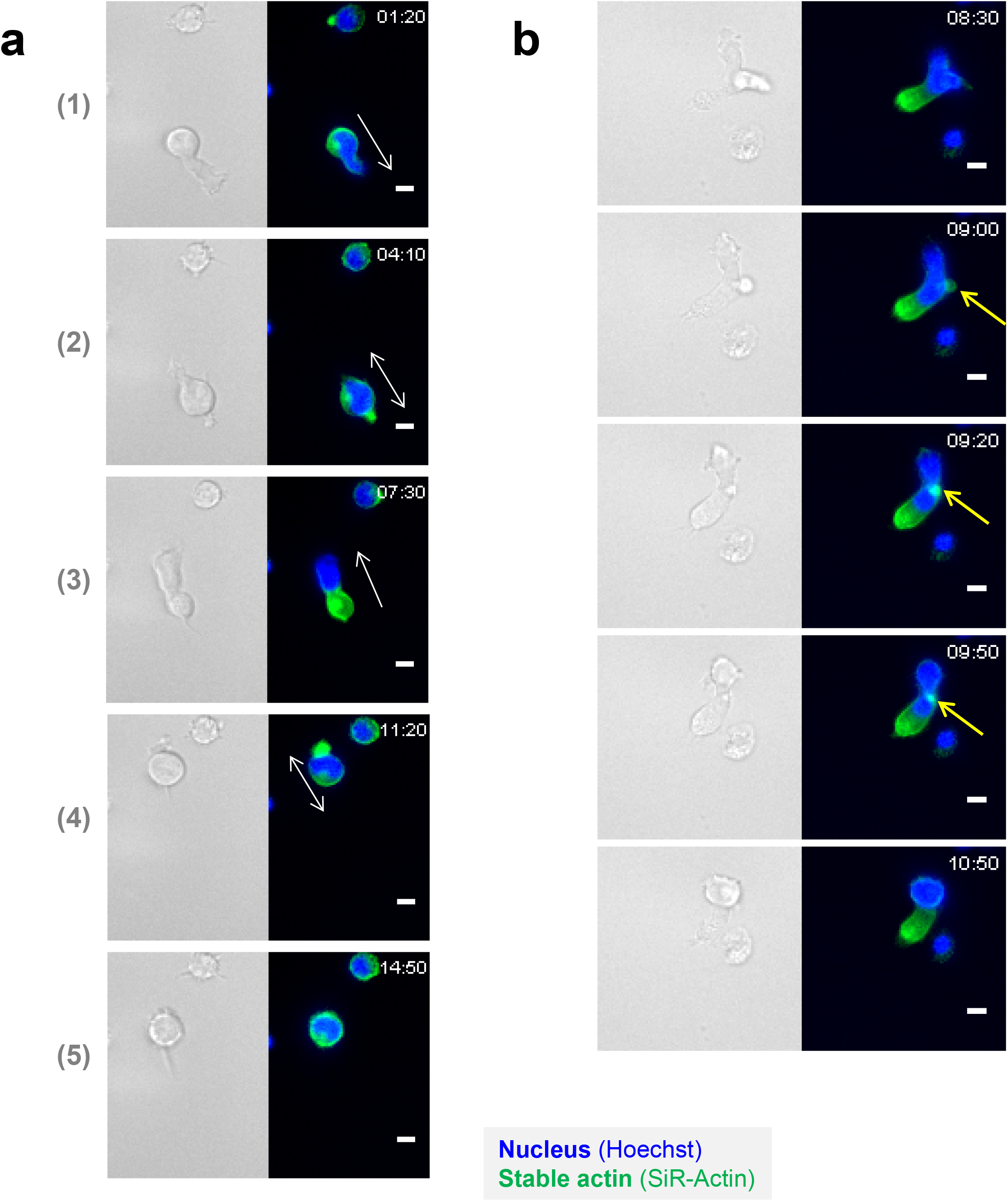
Stable actin relocalization upon lamellipodium retraction. Cells were labeled with Hoechst for the nucleus and SiRActin for the stable actin. Scale bar = 10μm **a**-Images of the migrating cell whose kymograph is presented in Fig 2a. **b**-Example of a cell developing two simultaneous lamellipodia. The local recruitment of stable actin (yellow arrow) allows the retraction of one of them.

**Supplementary Figure 3:**
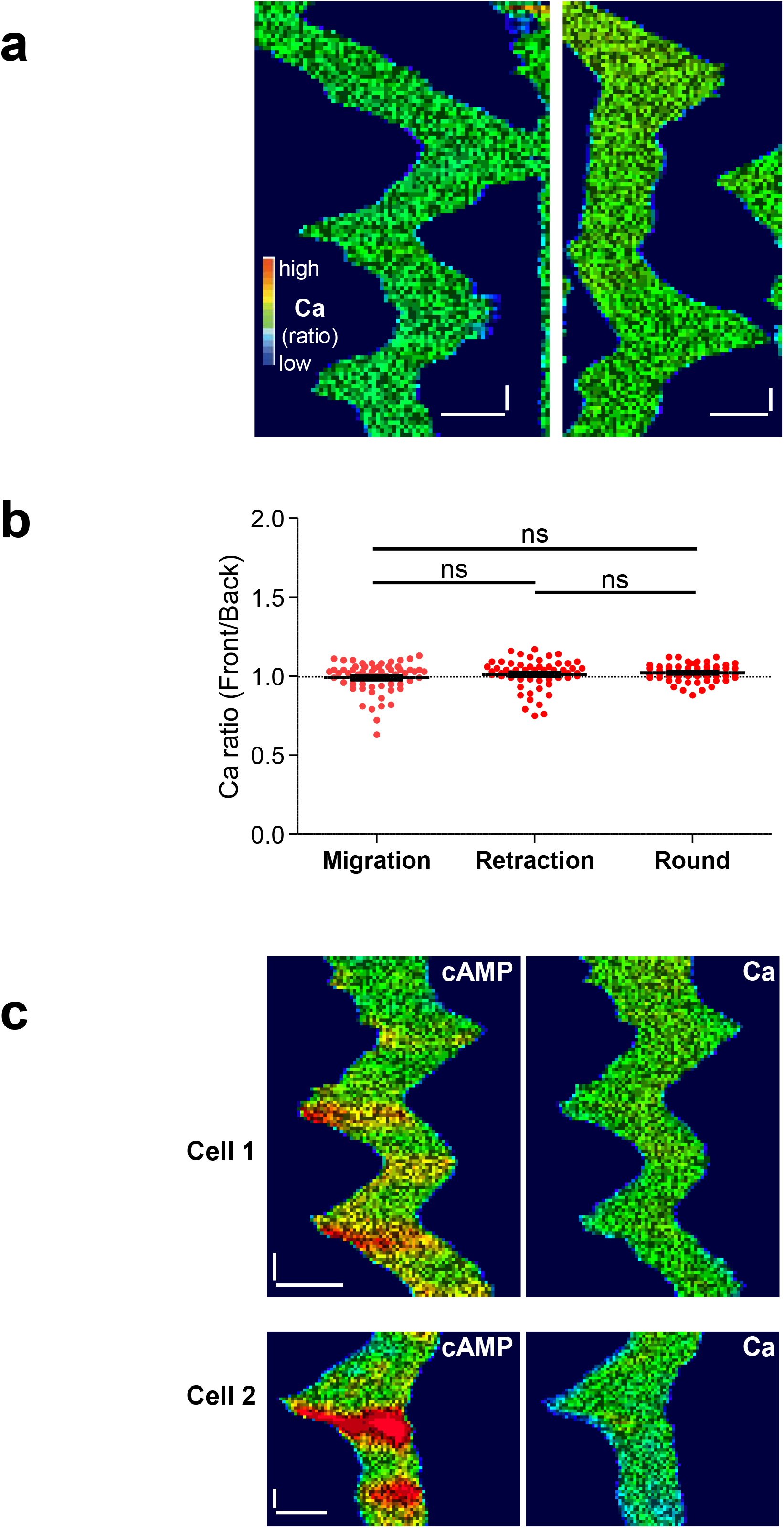
Calcium levels during migration. **a**-Kymographs of two examples of calcium measurements in Fura-2-loaded T cells deposited on CXCL12/VCAM-1 coated coverslips. No variation in Calcium could be detected. Calcium level is coded in false color. Horizontal scale bar = 10μm, vertical scale bar = 1min. **b**-Ratios between the front and the back levels of Calcium have been measured by drawing scanlines along the cell axis in cells migrating, retracting their lamellipodium or rounding up (after retraction). 57-59 values ± SE from 14 different cells. **c**-Kymographs of two examples of simultaneous cAMP (TEpacVV) and Calcium (Fura-2) recordings. Although some cAMP increase could be observed when cells changed direction, no calcium variations appeared. Horizontal scale bar = 10μm, vertical scale bar = 1min.

**Supplementary Figure 4:**
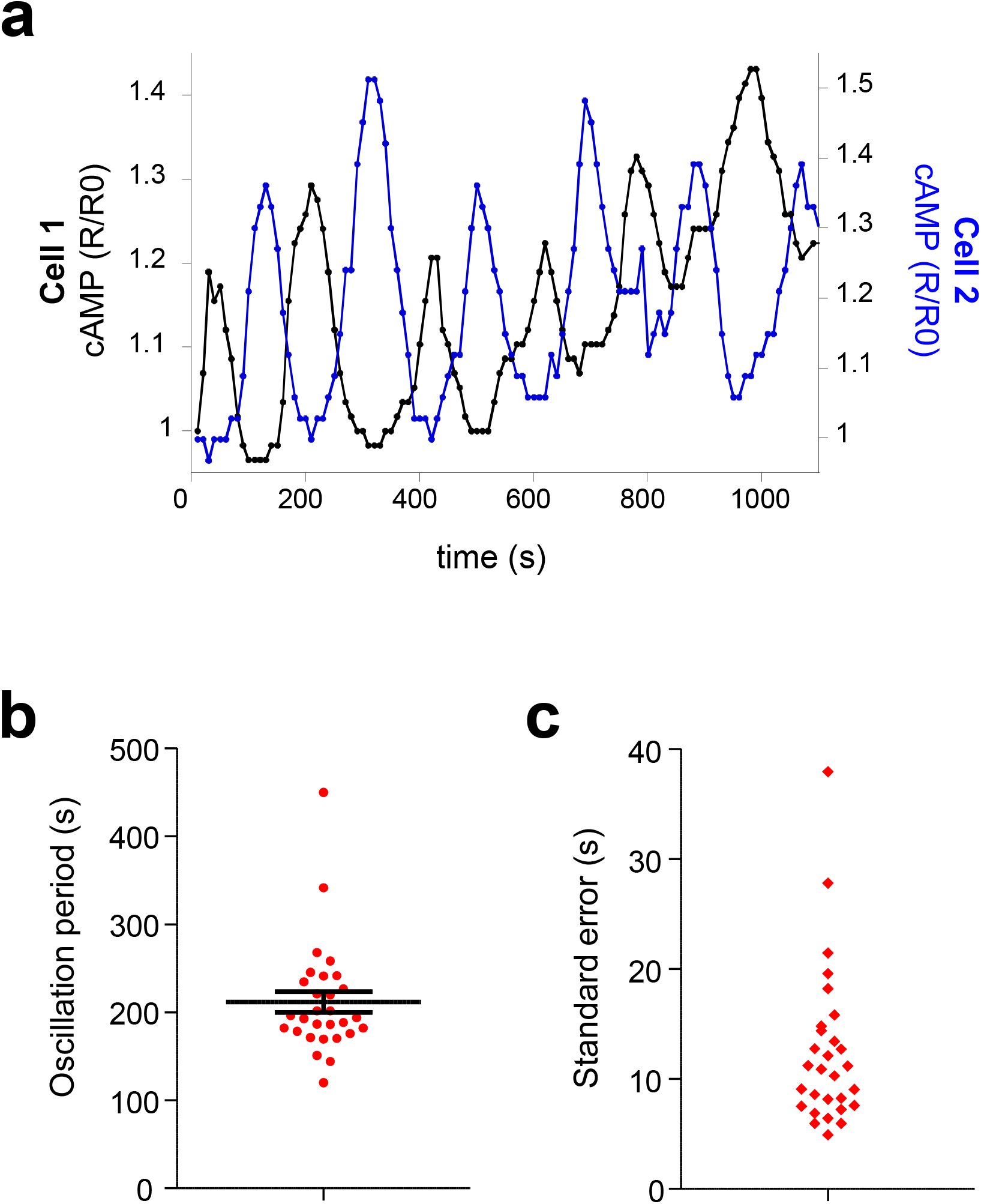
cAMP oscillations upon elongation/retraction cycles. **a**-Example of two different TEpacVV-transfected cells displaying cAMP oscillations. cAMP values have been normalized to the initial ratio (R0). **b**-Distribution of cAMP oscillating periods. The mean ± SE corresponds to the average period measured in 29 cells displaying 3-14 oscillations. Each dot represents one cell. Note the low dispersion of the period among the different cells. **c**-For each cell displaying 3-14 cAMP oscillations, the delay between 2 cAMP peaks was measured and the average period calculated. The regularity of the oscillations was assessed by calculating the standard error around their average. The small dispersion of the values indicates that, for a given cell, the oscillations are very regular.

**<u>Supplementary Movie 1</u>: Random migration on VCAM-1 / CXCL12 coated coverslip**

Transmitted light images of T lymphocytes migrating randomly. Scale bar = 10μm.

**<u>Supplementary Movie 2</u>: Asymmetry of actin networks upon chemokine-stimulated T cells**

Example of a T cell transfected with LifeAct-mCherry and labelled with SiRActin and Hoechst deposited on a VCAM-1/CXCL12-coated coverslip and observed. Image analysis of this cell in Fig 1a. Scale bar = 10μm.

**<u>Supplementary Movie 3</u>: Stable actin relocalization**

Example of a T lymphocyte labelled with SiRActin and Hoechst deposited on a VCAM-1/CXCL12-coated coverslip and monitored upon migration. The corresponding kymograph is shown in Fig 2a. Scale bar = 10μm.

**<u>Supplementary Movie 4</u>: cAMP variations upon migration**

TEpacVV-transfected T cell migrating on a CXCL12/VCAM-1-coated coverslip. cAMP is coded in false colors. The corresponding kymograph is shown in Fig 3a. Scale bar = 10μm.

**<u>Supplementary Movie 5</u>: cAMP oscillations upon elongation/retraction cycles**

TEpacVV-transfected T cell migrating on a CXCL12/VCAM-1 coated coverslip. cAMP is coded in false colors. The corresponding kymograph is shown in Fig 3d. Scale bar = 10μm.

**<u>Supplementary Movie 6</u>: cAMP increase and stable actin relocalization**

TEpacVV-transfected T cell loaded with SiRActin deposited on a CXCL12/VCAM-1-coated coverslip. Detailed images in Fig 5a. cAMP is coded in false colors. Scale bar = 10μm.

**<u>Supplementary Movie 7</u>: Effect of caged-cAMP photo-release on stable actin relocalization and cell rounding**

The cell was incubated previously in DMACM-caged 8-Br-cAMP and labeled with SiRActin. At 20s, it was illuminated at 405nm on a spot of 7μm diameter (white circle). Detailed images in Fig 4c. Scale bar = 10μm.

